# Ciprofloxacin-based ionic liquids demonstrate increased mutation frequency in *Escherichia coli*

**DOI:** 10.1101/2025.09.29.679178

**Authors:** Patrick Mikuni-Mester, Birgit Bromberger, Timea Dömök, Daniela Zetner, Laura Schleifer, Olga Makarova

**Affiliations:** Centre for Food Science and Veterinary Public Health, Unit of Food Microbiology, University of Veterinary Medicine, Vienna; Centre for Food Science and Veterinary Public Health, Unit of Veterinary Public Health and Epidemiology, University of Veterinary Medicine, Vienna

**Keywords:** ionic liquids, Escherichia coli, mutation frequency, antibiotics, API-ILs, ciprofloxacin

## Abstract

In the last decade, development of active pharmaceutical ingredient ionic liquids (API-ILs) has been proposed as a prospective game-changing strategy to overcome multiple problems with conventional solid-state drugs. Formulation of antibiotics as API-ILs has been reported to improve physico-chemical parameters as well as possible synergistic effects overcoming antimicrobial resistance. What has not been assessed to date however are possible adverse effects of the API-ILs considering the potentially wide-scale application and the inevitable release to the environment, as is the case with antibiotics and other popular medications. In this study, API-ILs based on the antibiotic ciprofloxacin were synthesized and investigated in terms of their antimicrobial activity and influence on mutation frequency of *Escherichia coli* MG1655. This is the first report that by formulating of antibiotics as API-ILs, bacterial mutation frequencies can be significantly increased, while structurally similar chloride ILs alone do not have such an effect. The observed changes in mutation frequencies seem to be dependent on the structure of the respective counterion. As increased mutation frequencies can fuel adaptation and ultimately resistance development of bacteria, it is crucial to improve our understanding of how API-ILs can designed in a safer way.

## 1. Introduction

Since their first introduction as novel “green” solvents and enhanced reaction media, ionic liquids (ILs), organic salts with melting points below 100°C, have found their way into a manifold of different applications covering the entire spectrum of the natural sciences. The increasing number of IL applications and their industrialization, particularly in organic synthesis, catalysis, extraction processes, electrochemistry and as active pharmaceutical ingredients has prompted investigation of their hazard potential in different biological test systems [1–9]. Their environmental fate has been studied in terms of persistence in air, water and soil, (bio)degradation, bioaccumulation in aquatic or terrestrial organisms and their (eco)toxicity, which is dependent on their specific structure, such as the head group or the side-chain of the cation or the anion [9–17]. While significant biological activity might be unfavorable in cases where ILs are utilized as solvents or catalysts, their toxicity can be considered valuable for pharmaceutical and medical uses. In specific contexts, the antimicrobial attributes of ILs can be harnessed, and their efficacy against antibiotic-resistant pathogens has already been demonstrated [2,5,18–22]. The combinatorial modification of the cation-anion composition and additional modification of the chemical structure of each ion allows adjustment of the expected toxicity of ILs towards a particular pathogen. Initially, it has been proposed that the primary mode of action of ILs, particularly those with elongated alkyl side chains, involves disrupting cell membrane function or even compromising cell integrity. However, it has been demonstrated that ILs can also engage with essential cellular elements such as proteins and DNA, inducing oxidative stress and disrupting metabolic functions [21,23]. The wide variety of cellular targets on the one hand might explain their effectivity against antimicrobial resistant (AMR) bacteria [13,21,24,25] but potentially could lead to increased development of new resistances due to increased mutation rates [26,27]. Surprisingly, this important aspect of ILs has not been experimentally tested so far.

In addition to “traditional” antimicrobial ILs, since 2007 the concept of using ILs as *pharmaceuticals* was developed and over the last 15 years numerous active pharmaceutical ingredient ionic liquids (API-ILs) have been described and studied [28,29]. The API-IL approach is able to overcome multiple problems with conventional solid state drugs, for example, bioavailability or polymorphism and drugs that have been prepared as API-ILs include Alzheimer’s and Parkinson’s disease medication, antibiotics, anticancer drugs, antibacterial and antifungal drugs, antihistamine and calcium channel blockers, numerous antimalarial APIs, antivirals, arthritis drugs, beta blockers, drugs for diabetes, gastrointestinal disorders, hyperkalemia, local anesthesia, pain relievers, multiple respiratory disorder medications, supplements, and vitamins, among others [28].

API-ILs offer the potential to, by adjusting the counterion, optimize the chemical and physical properties (e.g., hydrophilic−lipophilic balance (HLB), melting point, hydrophilicity and more) but also biological activity is a tunable property. One drug class where the biological activity of API-ILs has received significant attention are antibiotics and API-ILs based on ampicillin [24,30,31], nalidixic acid [25,32] and tetracycline [33] have been successfully synthesized and described. The focus so far was on improving either solubility, stability or bioavailability of these antibiotics or introduce a secondary mode of action to overcome bacterial resistance. Especially the latter would be a game changer as it would offer the potential to fight back on developing (and already developed) antibiotic resistances in bacterial pathogens, which is one of the main concerns of the WHO. Bacteria have developed a variety of adaptations over billions of years that can enhance their antimicrobial resistance, including expression of stress response genes and metabolic changes [34,35]. The ever-increasing spread of antibiotic and antimicrobial resistance creates the risk of a reversal to a world similar to before the advent of modern antibiotics.

Bacteria can be intrinsically resistant to certain antibiotics but also acquire antibiotic resistance by *de novo* mutations in chromosomal genes that can become mobilized and spread through horizontal gene transfer. Mutation rate, which is determined as the rate, at which spontaneous mutations arise per cell division and a closely related concept of mutation frequency (number of mutants present in a population at a given time point) play an important role in adaptive evolution under the selective pressure, such as that exerted by antimicrobials [36]. It has been previously shown that many antibiotics activate bacterial stress responses, leading to the expression of error-prone polymerases that result in increased mutation rates [37,38]. These increased mutation rates contribute to the increased standing genetic variation, on which selection can act, therefore creating the vicious cycle whereby antibiotic use fuels antibiotic resistance. Indeed, when treated with Ampicillin, Ciprofloxacin and Kanamycin, bacteria show 3 to 4 fold increase in mutation rate, which are closely linked with pathogen adaptation to the host [39] and evolutionary rescue of populations of microorganisms [40]. It is therefore a main goal to develop new antibiotics that do not lead to increases in mutation rates.

In this study, we investigated the mutation frequency of API-ILs based on a popular antibiotic that is on the WHO list of essential medicines – ciprofloxacin (fluoroquinolones). Ciprofloxacin is known to increase mutation rate of bacteria and previous studies already demonstrated that API-ILs with synergistic effects and improved bioavailability can be synthesized [41,42]. For this study API-ILs with five different counter cations were synthesized. The five counter cations comprised of two non-active, one medium active and two highly antimicrobially active structural motifs, which allows possible structure-dependent as well as activity-dependent effects if such effects exist. By also including “normal” chloride-based ILs of each counter cation, as well as the pure antibiotics, we also can determine if possible changes in mutation frequency are connected to the ILs per se or are the consequence of the API-IL. To the best of our knowledge, this study is the first study of possible effects of ionic liquids or API-ILs on bacterial mutation frequencies. Information on potential mutagenicity of API-ILs is not only important for potential application as novel antibiotics but also considering their increasing presence in natural environments [43].

## 2. MATERIALS AND METHODS

### 2.1. Ionic liquids and other chemical substances

The five chloride-based ionic liquids used in this study were provided by Proionic GmbH (Grambach, Austria) with a nominal purity of >98%. All API-ILs based on Ciprofloxacin were synthesized in our laboratory, according to the CBILS© route (CBILS is a registered trademark of Proionic GmbH) [44,45]. Cationic precursors for API-IL synthesis were provided by Proionic GmbH (Grambach, Austria) with a nominal purity of >98% and ciprofloxacin was purchased from Sigma-Aldrich (Wien, Austria). All investigated ILs and API-ILs are summarized in table 1 and Figure 1.

**Table 1.**
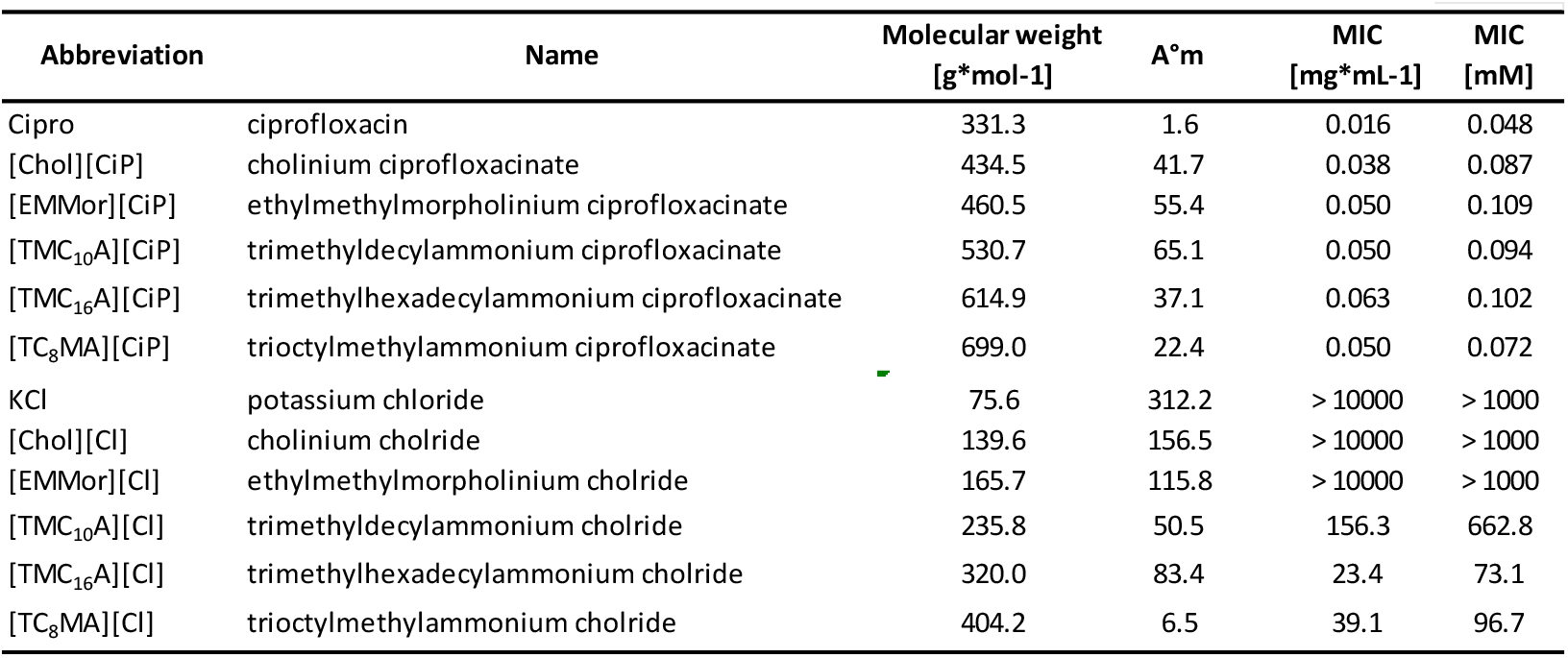
List of all investigated ILs, API-ILs and antibiotics including name, abbreviation, molecular weight, limiting molar conductivity and MIC (mg/L and mM).

| Abbreviation | Name | Molecular weight<br>[g*mol<sup>-1</sup>] | A°m | MIC<br>[mg*mL<sup>-1</sup>] | MIC<br>[mM] |
| --- | --- | --- | --- | --- | --- |
| Cipro | ciprofloxacin | 331.3 | 1.6 | 0.016 | 0.048 |
| [Chol][CiP] | cholinium ciprofloxacin | 434.5 | 41.7 | 0.038 | 0.087 |
| [EMMor][CiP] | ethylmethylmorpholinium ciprofloxacin | 460.5 | 55.4 | 0.050 | 0.109 |
| [TMC<sub>10</sub>A][CiP] | trimethyldecylammonium ciprofloxacin | 530.7 | 65.1 | 0.050 | 0.094 |
| [TMC<sub>16</sub>A][CiP] | trimethylhexadecylammonium ciprofloxacin | 614.9 | 37.1 | 0.063 | 0.102 |
| [TC<sub>8</sub>MA][CiP] | trioctylmethylammonium ciprofloxacin | 699.0 | 22.4 | 0.050 | 0.072 |
| KCl | potassium chloride | 75.6 | 312.2 | > 10000 | > 1000 |
| [Chol][Cl] | cholinium chloride | 139.6 | 156.5 | > 10000 | > 1000 |
| [EMMor][Cl] | ethylmethylmorpholinium chloride | 165.7 | 115.8 | > 10000 | > 1000 |
| [TMC<sub>10</sub>A][Cl] | trimethyldecylammonium chloride | 235.8 | 50.5 | 156.3 | 662.8 |
| [TMC<sub>16</sub>A][Cl] | trimethylhexadecylammonium chloride | 320.0 | 83.4 | 23.4 | 73.1 |
| [TC<sub>8</sub>MA][Cl] | trioctylmethylammonium chloride | 404.2 | 6.5 | 39.1 | 96.7 |

**Fig. 1:**
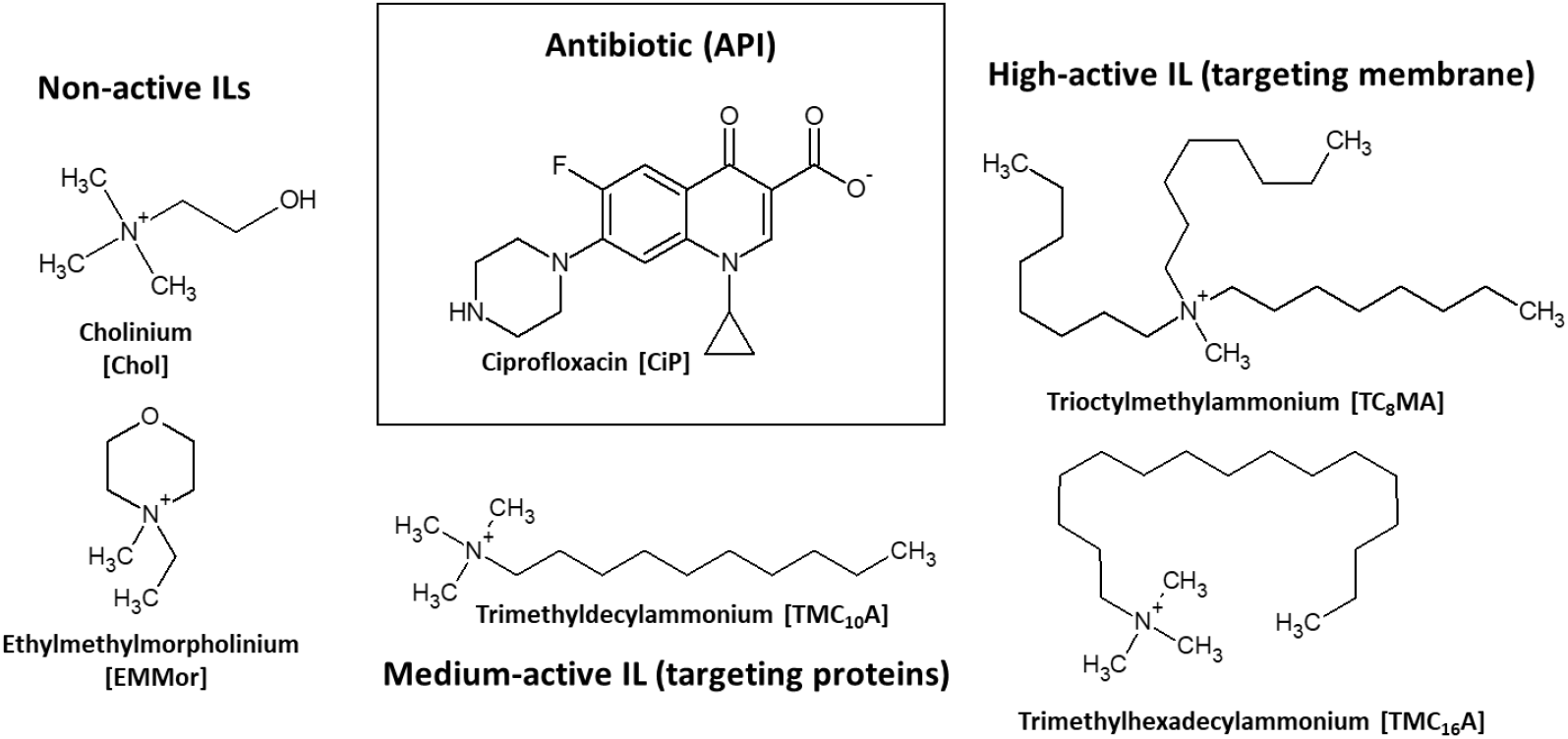
Schematic representation of the respective cations and anions of the 5 API-ILs investigated in this study.

### 2.2. Conductivity measurements

The specific conductivities were measured using a Seven Compact Cond meter S230 (Mettler Toledo). Initially, 10 ml of a 25 mM aqueous solution (ddH_2_O) of each compound was prepared and conductivity measurements were made at 25°C. A known amount of ddH_2_O was added stepwise to the sample to obtain the required concentration (50 mM to 1.5 mM) and conductivity was measured after each addition (see tables S1&2). The molar conductivity was calculated using equation Λ_m_ = (k/c). The limiting molar conductivity Λ°_m_ was determined graphically by plotting 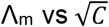 according to Kohlrausch square root law 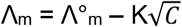.

### 2.3. Bacterial strains and culture conditions

*Escherichia coli* K-12 MG1655 was used for all experiments, grown overnight at 37°C in LB broth (Oxoid™, Hamsphire, United Kingdom). Bacteria were maintained at −80 °C using Microbank™ technology (Pro-Lab Diagnostics, Richmond Hill, Canada).

### 2.4. Minimal inhibitory concentration (MIC) assessment and disc diffusion

MICs of the test chemicals (ILs, API-ILs and ciprofloxacin) were assessed by applying the serial two-fold dilution microtiter plate method in LB broth [46]. In order to create a constant cell status for each experiment, 1 ml aliquots of the respective overnight cultures were transferred into 9 ml of fresh LB medium (1:10 dilution) and incubated for 3 h at 37 °C to ensure that cells were in a logarithmic growth-phase. Subsequently, each well, which contained a serial diluted antimicrobial substance (dilution 1:2), was inoculated with 5 × 10^5^ CFU bacterial cells. After inoculation, absorbance of the 96-well microtiter plates (Corning B.V Life Sciences, Amsterdam, Netherlands) was measured at a wavelength of 610 nm in a TECAN F100 microplate reader (Tecan Austria GmbH, Groeding, Austria) to monitor for any possible interference by the antimicrobial substances. The microtiter plates were then incubated for 24 h at the 37°C and bacterial growth assessed visually and confirmed by measuring the absorbances at 610 nm. The MIC was defined as the lowest concentration of the tested antimicrobial substance where no bacterial growth could be measured after 24 h. Results are presented as mean MICs and at least two experiments were performed on different days. Each experiment included positive (bacterial growth control in LB) and negative controls (medium without the addition of bacteria).

For the diffusion test with *E. coli*, 1 ml of an overnight culture was used to inoculate 9 ml of TSB. This culture was grown for three hours with vigorous shaking at 37 °C to ensure that the cells were in the logarithmic growth phase. The suspension was diluted with TSB to give a respective 5 × 10^5^ CFU and immediately swabbed onto TSA plates. Up to six paper filter disks of 6 mm in diameter were aseptically placed on each plate. Afterwards, ten microliters of each IL, API-IL and ciprofloxacin dilution were applied per disc. The plates were incubated at 37 °C for 18 hours. Finally, the circular zones of inhibition were measured with a ruler.

### 2.5 Mutagenesis experiments

To determine mutagenesis, the mutation frequency was determined as described previously [47]. To obtain comparable results from the activity of classic antibiotics, ILs and API-ILs, the mutation frequency experiments were performed as follows. Three ml of an overnight culture *E. coli* MG1655 were diluted 1:100 in fresh LB broth and incubated for two hours at 37°C at 200 rpm until an OD_600nm_ of 0.5 was reached. Afterwards the subculture of *E. coli* MG1655 was again diluted 1:100 in LB medium containing ½ MIC concentration of the test compound and incubated overnight at 37°C and 200 rpm. After incubation, the bacterial cells were pelleted via centrifugation (4.000*g for 10 min) and washed twice with 0.9% saline solution. After the final washing step, cells were resuspended in 10 ml fresh LB broth and incubated overnight at 37°C and 200 rpm. After incubation, bacterial concentrations were determined by spot plating on LB agar plates and the CFU/ml calculated. To determine the number of mutants, 10 ml of the bacterial culture was pellet via centrifugation, resuspended in 1 ml of 0.9% saline solution and 100µl plated on LB agar plates containing 100 µg/ml Rifampicin. Mutation frequencies were calculated by maximum verisimilitude method and data were processed using the on-line web-tool for mutation frequency determination Falcor http://www.mitochondria.org/protocols/FALCOR.html). Each compound was tested on five different days with five independent replicates.

## 3. Results and Discussion

### 3.1. Characterization of antimicrobial activity of novel API-ILs

In this study, novel API-ILs based on the antibiotic ciprofloxacin were synthesized using the CBILs route, which is a completely halide and waste free process producing high-purity ionic liquids [48]. We chose a set of 5 counter ions with varying antimicrobial activity to investigate possible synergistic effects of the new API-ILs in terms of antimicrobial activity and mutation frequency. Cations [Chol]^+^ and [EMMor]^+^ have a low antimicrobial activity, [TMC_10_A]^+^ an intermediate and both [TMC_16_A]^+^ and [TC_8_MA]^+^ are highly antimicrobial due to the presence of long or multiple alkyl side chains. Minimal inhibitory concentration (MIC) for each antibiotic, the respective chloride ILs of the cations and the API-ILs were determined (Table 1). To account for the differences in molecular weight of the API-ILs, in addition to the unit mg/L, which is usually used for antibiotics, the respective mM concentrations are also given.

To determine the degree of API-IL dissociation in aqueous solution, conductivity for all compounds were performed and expressed as limiting molar conductivity (See supplemental tables S1&2 for full results). The degree of dissociation of API-ILs can support the identification of possible synergistic or sum effects of the respective cations and anions. Overall, a lower dissociation of the API-ILs compared to the respective chloride ILs was observed, indicating formation of stable ion pairs between the respective cation and anion.

For ciprofloxacin-based API-ILs, all MIC values are within the 2-fold resolution limit of the microbroth dilution method indicating a similar activity compared to pure ciprofloxacin. As the MIC values for the 5 API-ILs are significantly lower compared to the respective chloride ILs, no indication of an inactivation of the antibiotic during synthesis was found.

In addition to the microbroth dilution assay, the antimicrobial activity of the newly synthesized API-ILs, and respective chloride ILs, was additionally tested by disc diffusion test. For each compound three different concentrations were tested, and representative results are listed in table 2 (for complete results see supplemental Table S3). In case of the chloride ILs, only for [TMC_10_A][Cl] a small inhibition zone was detectable at 50 mg/disc, which was the highest tested concentration of the API-ILs.

**Table 2.** Mean inhibition zones [mm] and standard deviation of all 5 API-ILs, 5 ILs and pure antibiotic.

| API-IL - cation | Ciprofloxacin<br>(0.5 mg/disc) |  | chloride<br>(50 mg/disc) |  |
| --- | --- | --- | --- | --- |
|  | Mean | SD | Mean | SD |
| Pure antibiotic | 25.00 | 0.0 | - |  |
| [Chol] | 21.50 | 2.5 | n.d. |  |
| [EMMor] | 20.50 | 0.5 | n.d. |  |
| [TMC<sub>10</sub>A] | 20.50 | 3.5 | 8.5 | 0.5 |
| [TMC<sub>16</sub>A] | 21.00 | 4.0 | n.d. |  |
| [TC<sub>8</sub>MA] | 18.50 | 2.5 | n.d. |  |

Only small differences between the five API-ILs were detectable with [TC_8_MA][CiP] showing the smallest inhibition zone. However, inhibition zones for API-ILs were all smaller compared to pure ciprofloxacin. This effect can be due to the higher molecular weight of the API-ILs resulting in a lower amount of API molecules, but also slower diffusion rates of the API-IL or the [CiP]^-^ anion within the agar. A similar effect has been reported previously and thus it is recommendable to combine different methods when determining API-IL activity [49,50]. Overall, disc diffusion together with the MIC results confirms that the ciprofloxacin-based API-ILs demonstrated an equal antimicrobial activity as the pure ciprofloxacin.

In summary, the results of both the microbroth dilution assay and the disc diffusion assay confirm the successful synthesis of 5 API-ILs based on ciprofloxacin. The API-ILs showed activity both in the microbroth dilution test as well as the disc diffusion test, with the latter revealing different diffusion rates of the API-ILs compared to the pure antibiotics. This effect could be either by the bigger size and increased hydrophobicity of the respective associated API-ILs anions, which would explain why the bulky [TC_8_MA]^+^ based API-IL demonstrated the smallest inhibition zones.

### 3.2 Mutagenesis of API-ILs

For analysis of API-IL mutagenesis, the ½ MIC concentrations was used for each of the API-ILs, pure ciprofloxacin and the five chloride ILs. This concentration was chosen based on own preliminary and published data demonstrating that half MIC induced the highest increase in mutagenesis by ciprofloxacin [51], and the expectation that environmental contamination would likely be at the sub-MIC concentration due to the dilution effects. To ensure that bacteria were still able to grow under the given concentrations, growth curves were performed for each substance (figure 1).

**Fig. 1:**
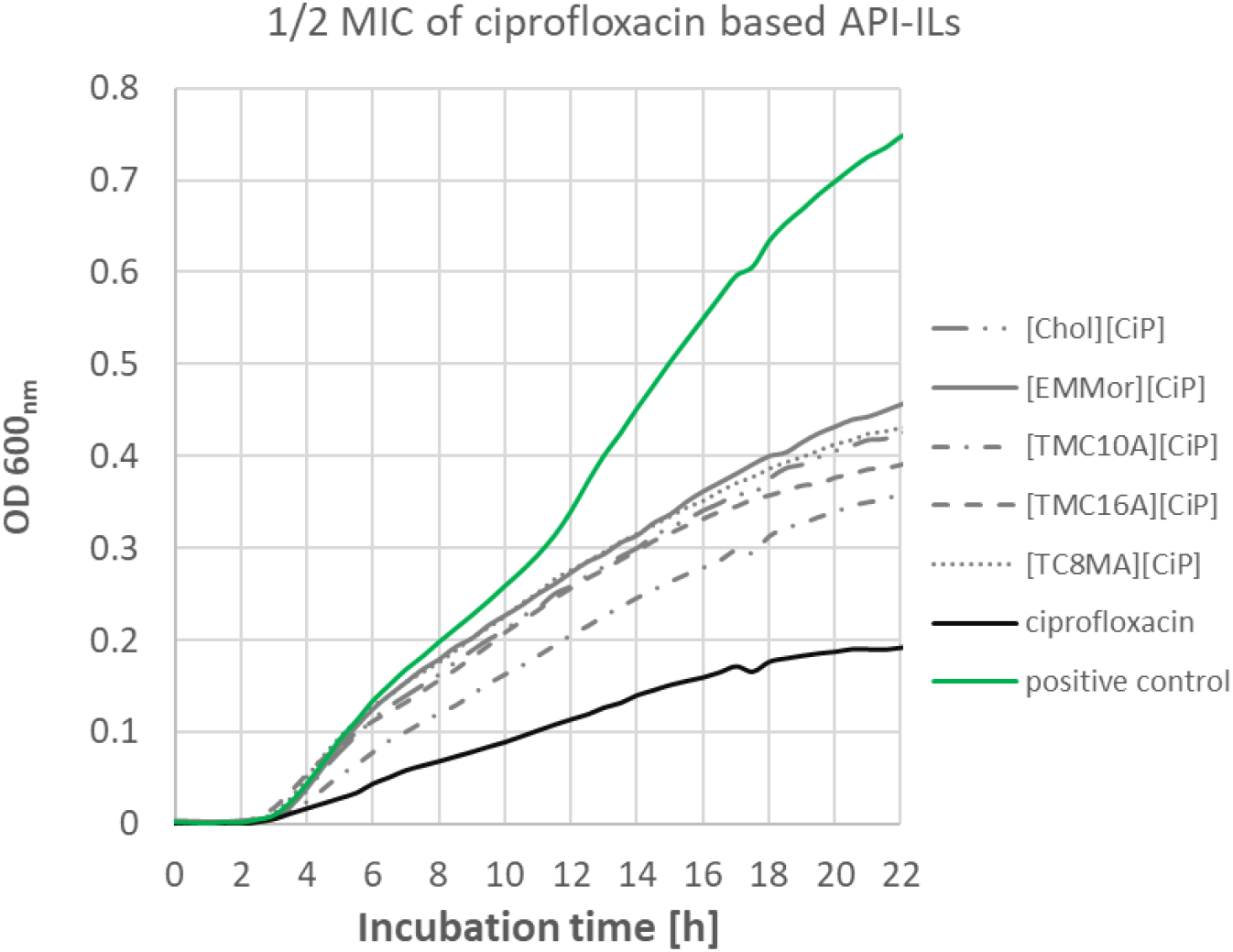
Growth curves of *E. coli* MG1655 at ½ MIC of 5 API-ILs and pure antibiotics.

Figures 2 and 3 depict the median mutation frequency for *E. coli* MG 1655 exposed to all substances shown (see table S5). In good accordance with the literature, higher mutation frequencies were found for bacteria exposed to ciprofloxacin, although the increase of 1.71 was lower compared to previous studies in which 3 to 4-fold increases were detected [52]. The discrepancy can be due to differences in experimental conditions, such as different concentrations of the drugs and duration of treatment that have been shown to affect mutation rate [36]. For the chloride ILs, only in the case of [TMC_10_A][Cl] a 1.6 fold higher mutation frequency was determined, while for the other chloride ILs no impact on mutation frequency was found compared to the non-treatment controls (Figure 3). Therefore a possible impact of the respective cations on mutation frequency changes in the API-ILs can be excluded, with the exception of those with the [TMC_10_A]^+^ cation. It is important to point out that we did evaluate the possible mutagenesis of ILs per se, as only relatively low concentrations based on the respective API-ILs were tested. To the best of our knowledge, to date solely directed evolution experiments to increase ionic liquid tolerance (mostly for biomass applications) have been conducted but nothing is known regarding the general or a structure dependent mutagenicity of ionic liquids [53,54].

**Fig. 2:**
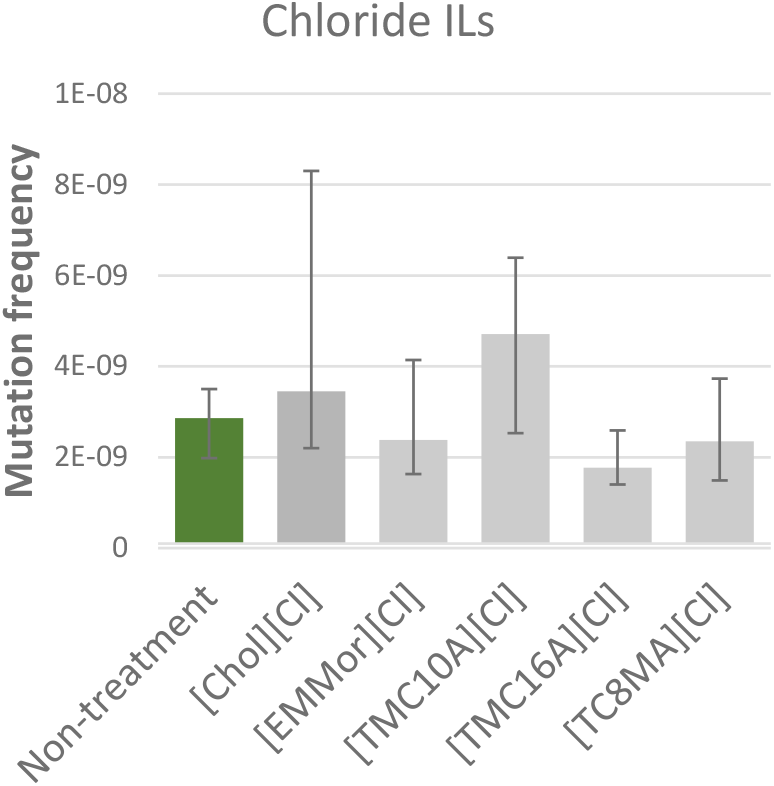
Changes in mutation frequency in *E. coli MG1655* induced by chloride based ionic liquids at ½ MIC. Error bars represent 95% confidence interval for mutation frequency estimation by plating in Rifampicin (100 mg/ml) and using the maximum likelihood method.

**Fig. 3:**
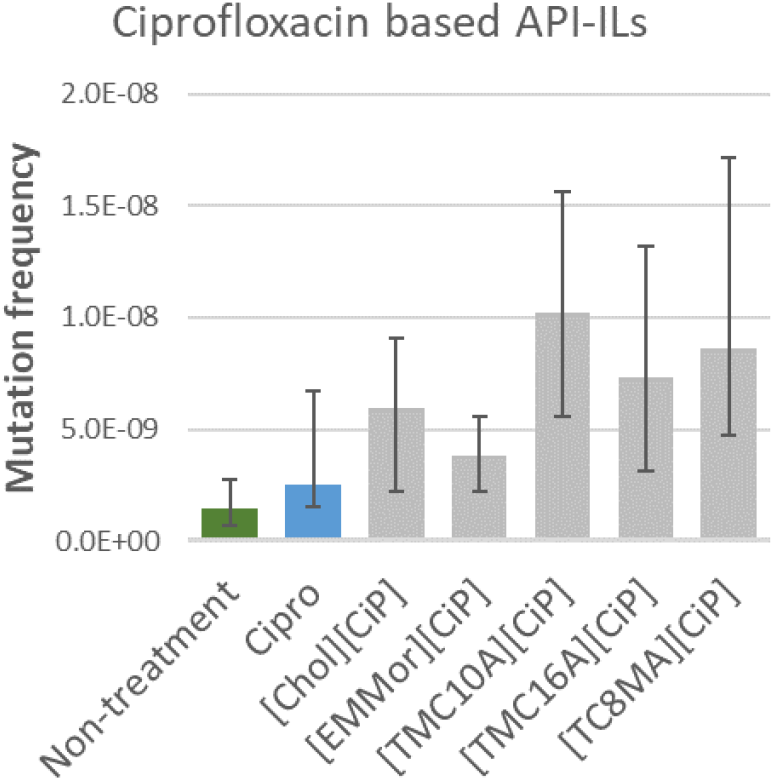
Changes in mutation frequency in *E. coli MG1655* induced by ciprofloxacin-based API-ILs at ½ MIC. Error bars represent 95% confidence interval for mutation frequency estimation by plating in Rifampicin (100 mg/ml) and using the maximum likelihood method.

For ciprofloxacin-based API-ILs, not only were the observed mutation frequencies higher compared to the non-treatment control but also compared to ciprofloxacin alone. While for [EMMor][CiP] the mutation frequency was only slightly increased, for the other API-ILs a 2-3-fold higher mutation frequency compared to ciprofloxacin could be found (Figure 3). Given the fact that, due to the higher molecular weight, the molecular equivalent of ciprofloxacin is lower, a clear connection between the formulation as an API-IL to an increased mutation frequency is clear. As the respective chloride ILs did not lead to a similar increase at significantly higher concentrations, it is unlikely that the observed increased mutagenesis is a simple sum of the anion (in this case ciprofloxacin) and cation effects but a new intrinsic property of the API-IL. However, we also found clear differences between the API-ILs with non-toxic to bacteria cations ([Chol][Cip] and [EMMor][Cip]) and the antimicrobially active ones ([TMC10A][Cip], [TMC16A][Cip] and [TC8MA][Cip], demonstrating once more that the respective molecule structure can influence the outcome. As all the three active cations target or at least interact with bacterial membranes, an increased penetration of the API-IL or reduced efflux could be the underlying cause. It will be important in future studies to further increase the structural diversity of API-IL cations to include different head groups, side chain functionalization but also different antimicrobial modes of action [1,21]. The ultimate goal would be to develop quantitative structure–activity relationship (QSAR) models for mutation frequency prediction that can serve as an early warning system for mutagenicity of novel API-ILs, not only for procaryotic but also eucaryotic cells.

## 4. Conclusions

This is the first report demonstrating that changing the phyisco-chemical and biological properties of pharmaceuticals (in this case antibiotics) by transforming them into API-ILs can significantly influence bacterial mutation frequency, while the respective chloride ILs did not. By combining ciprofloxacin with 5 counter ions of different antimicrobial activity, we could demonstrate that this effect is not a general aspect of ionic liquids or even API-ILs *per se*, but structure dependent. Nevertheless, increased mutation frequency was found for all API-ILs demonstrating a higher mutagenesis of these compounds. As mutation frequency is generally connected to bacteria’s possibility to develop antimicrobial resistance, this seems to be a potential drawback to the API-IL approach that needs to be investigated further. Future studies, such as experimental evolution, can help establish whether the increased mutation frequency indeed contributes to an increased resistance to antimicrobials and whether this relationship is structure-dependent. Altogether, this will greatly contribute to a better development of API-ILs with a safer resistance profile as well as help to identify potential risks in resistance induction potential of the new IL-based antibiotics.

## CRediT authorship contribution statement

**Patrick Mikuni-Mester, Olga Makarova**: Conceptualization, Methodology, Writing, Discussion, Data treatment, Supervision. **Birgit Bromberger**: API-IL synthesis, Experiments, Writing. **Timea Dömök, Daniela Zetner**: Conductivity measurements, Mutagenesis experiments and MIC testing. **Laura Schleifer**: Mutagenesis experiments and MIC testing.

## Acknowledgments

This project received funding from the Department 3 of the University of Veterinary medicine, Vienna.

## Appendix A. Supporting information

Supplementary data associated with this article can be found in the online version at

