## Supplemental Material for "Ciprofloxacin-based ionic liquids demonstrate increased mutation frequency in *Escherichia coli*"

**Table S1:** Conductivity measurements of ciprofloxacin-based API-ILs and pure ciprofloxacin.

| | Concentration (mol) | √C | Conductivity (mS) | Am | | Concentration (mol) | √C | Conductivity (mS) | Am |
|---|---|---|---|---|---|---|---|---|---|
| Cipro | 0.0250 | 0.16 | 0.007 | 0.3 | [$TMC_{10}A$] [CiP] | 0.0250 | 0.16 | 0.687 | 27.5 |
| | 0.0167 | 0.13 | 0.005 | 0.3 | | 0.0167 | 0.13 | 0.620 | 37.2 |
| | 0.0125 | 0.11 | 0.005 | 0.4 | | 0.0125 | 0.11 | 0.509 | 40.7 |
| | 0.0083 | 0.09 | 0.004 | 0.4 | | 0.0083 | 0.09 | 0.383 | 46.0 |
| | 0.0063 | 0.08 | 0.004 | 0.6 | | 0.0063 | 0.08 | 0.303 | 48.5 |
| | 0.0050 | 0.07 | 0.003 | 0.7 | | 0.0050 | 0.07 | 0.235 | 47.0 |
| | 0.0042 | 0.06 | 0.003 | 0.8 | | 0.0042 | 0.06 | 0.209 | 50.2 |
| | 0.0036 | 0.06 | 0.003 | 0.8 | | 0.0036 | 0.06 | 0.183 | 51.2 |
| | 0.0031 | 0.06 | 0.003 | 0.9 | | 0.0031 | 0.06 | 0.165 | 52.7 |
| | 0.0025 | 0.05 | 0.003 | 1.1 | | 0.0025 | 0.05 | 0.137 | 54.6 |
| | 0.0021 | 0.05 | 0.003 | 1.2 | | 0.0021 | 0.05 | 0.107 | 51.4 |
| | 0.0018 | 0.04 | 0.002 | 1.4 | | 0.0018 | 0.04 | 0.100 | 56.1 |
| | 0.0016 | 0.04 | 0.003 | 1.7 | | 0.0016 | 0.04 | 0.090 | 57.5 |
| [Chol] [CiP] | 0.0250 | 0.16 | 0.638 | 25.5 | [$TMC_{16}A$] [CiP] | 0.0250 | 0.16 | 0.174 | 6.9 |
| | 0.0167 | 0.13 | 0.626 | 37.6 | | 0.0167 | 0.13 | 0.187 | 11.2 |
| | 0.0125 | 0.11 | 0.459 | 36.7 | | 0.0125 | 0.11 | 0.117 | 9.3 |
| | 0.0083 | 0.09 | 0.320 | 38.4 | | 0.0083 | 0.09 | 0.155 | 18.6 |
| | 0.0063 | 0.08 | 0.246 | 39.4 | | 0.0063 | 0.08 | 0.120 | 19.3 |
| | 0.0050 | 0.07 | 0.190 | 38.0 | | 0.0050 | 0.07 | 0.100 | 20.0 |
| | 0.0042 | 0.06 | 0.159 | 38.2 | | 0.0042 | 0.06 | 0.087 | 20.9 |
| | 0.0036 | 0.06 | 0.127 | 35.7 | | 0.0036 | 0.06 | 0.079 | 22.1 |
| | 0.0031 | 0.06 | 0.120 | 38.5 | | 0.0031 | 0.06 | 0.075 | 23.9 |
| | 0.0025 | 0.05 | 0.093 | 37.1 | | 0.0025 | 0.05 | 0.063 | 25.3 |
| | 0.0021 | 0.05 | 0.080 | 38.2 | | 0.0021 | 0.05 | 0.058 | 27.7 |
| | 0.0018 | 0.04 | 0.066 | 37.2 | | 0.0018 | 0.04 | 0.052 | 28.9 |
| | 0.0016 | 0.04 | 0.060 | 38.4 | | 0.0016 | 0.04 | 0.048 | 31.0 |
| [EMMor] [CiP] | 0.0250 | 0.16 | 1.071 | 42.8 | [$TC_8MA$] [CiP] | 0.0250 | 0.16 | 0.148 | 5.9 |
| | 0.0167 | 0.13 | 0.982 | 58.9 | | 0.0167 | 0.13 | 0.149 | 8.9 |
| | 0.0125 | 0.11 | 0.726 | 58.1 | | 0.0125 | 0.11 | 0.109 | 8.8 |
| | 0.0083 | 0.09 | 0.462 | 55.4 | | 0.0083 | 0.09 | 0.106 | 12.7 |
| | 0.0063 | 0.08 | 0.342 | 54.7 | | 0.0063 | 0.08 | 0.071 | 11.3 |
| | 0.0050 | 0.07 | 0.267 | 53.4 | | 0.0050 | 0.07 | 0.059 | 11.8 |
| | 0.0042 | 0.06 | 0.220 | 52.8 | | 0.0042 | 0.06 | 0.062 | 15.0 |
| | 0.0036 | 0.06 | 0.194 | 54.3 | | 0.0036 | 0.06 | 0.059 | 16.6 |
| | 0.0031 | 0.06 | 0.167 | 53.4 | | 0.0031 | 0.06 | 0.053 | 16.9 |
| | 0.0025 | 0.05 | 0.134 | 53.6 | | 0.0025 | 0.05 | 0.044 | 17.5 |
| | 0.0021 | 0.05 | 0.111 | 53.2 | | 0.0021 | 0.05 | 0.037 | 17.7 |
| | 0.0018 | 0.04 | 0.095 | 53.4 | | 0.0018 | 0.04 | 0.033 | 18.4 |
| | 0.0016 | 0.04 | 0.083 | 53.0 | | 0.0016 | 0.04 | 0.030 | 18.9 |

**Table S2:** Conductivity measurements of chloride ILs.

| | Concentration (mol) | √C | Conductivity (mS) | Am | | Concentration (mol) | √C | Conductivity (mS) | Am |
|---|---|---|---|---|---|---|---|---|---|
| KCl | 0.0250 | 0.16 | 3.320 | 132.8 | $[TMC_{10}A]$ [Cl] | 0.0250 | 0.16 | 0.567 | 22.7 |
| | 0.0167 | 0.13 | 2.650 | 159.0 | | 0.0167 | 0.13 | 0.717 | 43.0 |
| | 0.0125 | 0.11 | 2.330 | 186.4 | | 0.0125 | 0.11 | 0.549 | 43.9 |
| | 0.0083 | 0.09 | 1.074 | 128.9 | | 0.0083 | 0.09 | 0.384 | 46.1 |
| | 0.0063 | 0.08 | 1.416 | 226.6 | | 0.0063 | 0.08 | 0.287 | 45.9 |
| | 0.0050 | 0.07 | 1.197 | 239.4 | | 0.0050 | 0.07 | 0.241 | 48.2 |
| | 0.0042 | 0.06 | 1.024 | 245.8 | | 0.0042 | 0.06 | 0.185 | 44.4 |
| | 0.0036 | 0.06 | 0.888 | 248.6 | | 0.0036 | 0.06 | 0.163 | 45.7 |
| | 0.0031 | 0.06 | 0.777 | 248.6 | | 0.0031 | 0.06 | 0.152 | 48.6 |
| | 0.0025 | 0.05 | 0.639 | 255.6 | | 0.0025 | 0.05 | 0.124 | 49.6 |
| | 0.0021 | 0.05 | 0.538 | 258.2 | | 0.0021 | 0.05 | 0.092 | 43.9 |
| | 0.0018 | 0.04 | 0.465 | 260.4 | | 0.0018 | 0.04 | 0.089 | 49.6 |
| | 0.0016 | 0.04 | 0.407 | 260.5 | | 0.0016 | 0.04 | 0.077 | 49.1 |
| [Chol] [Cl] | 0.0250 | 0.16 | 2.990 | 119.6 | $[TMC_{16}A]$ [Cl] | 0.0250 | 0.16 | 0.469 | 18.8 |
| | 0.0167 | 0.13 | 1.988 | 119.3 | | 0.0167 | 0.13 | 0.716 | 43.0 |
| | 0.0125 | 0.11 | 1.536 | 122.9 | | 0.0125 | 0.11 | 0.528 | 42.2 |
| | 0.0083 | 0.09 | 1.255 | 150.6 | | 0.0083 | 0.09 | 0.462 | 55.4 |
| | 0.0063 | 0.08 | 0.915 | 146.4 | | 0.0063 | 0.08 | 0.349 | 55.8 |
| | 0.0050 | 0.07 | 0.729 | 145.8 | | 0.0050 | 0.07 | 0.301 | 60.2 |
| | 0.0042 | 0.06 | 0.635 | 152.4 | | 0.0042 | 0.06 | 0.257 | 61.7 |
| | 0.0036 | 0.06 | 0.534 | 149.5 | | 0.0036 | 0.06 | 0.241 | 67.5 |
| | 0.0031 | 0.06 | 0.477 | 152.6 | | 0.0031 | 0.06 | 0.210 | 67.2 |
| | 0.0025 | 0.05 | 0.369 | 147.6 | | 0.0025 | 0.05 | 0.175 | 69.8 |
| | 0.0021 | 0.05 | 0.315 | 151.2 | | 0.0021 | 0.05 | 0.141 | 67.5 |
| | 0.0018 | 0.04 | 0.268 | 150.1 | | 0.0018 | 0.04 | 0.123 | 69.1 |
| | 0.0016 | 0.04 | 0.233 | 149.1 | | 0.0016 | 0.04 | 0.103 | 66.1 |
| [EMMor] [Cl] | 0.0250 | 0.16 | 2.080 | 83.2 | $[TC_8MA]$ [Cl] | 0.0250 | 0.16 | 0.056 | 2.2 |
| | 0.0167 | 0.13 | 1.634 | 98.0 | | 0.0167 | 0.13 | 0.038 | 2.3 |
| | 0.0125 | 0.11 | 1.269 | 101.5 | | 0.0125 | 0.11 | 0.082 | 6.6 |
| | 0.0083 | 0.09 | 0.871 | 104.5 | | 0.0083 | 0.09 | 0.056 | 6.8 |
| | 0.0063 | 0.08 | 0.667 | 106.7 | | 0.0063 | 0.08 | 0.042 | 6.7 |
| | 0.0050 | 0.07 | 0.531 | 106.2 | | 0.0050 | 0.07 | 0.037 | 7.3 |
| | 0.0042 | 0.06 | 0.445 | 106.8 | | 0.0042 | 0.06 | 0.030 | 7.2 |
| | 0.0036 | 0.06 | 0.380 | 106.4 | | 0.0036 | 0.06 | 0.021 | 5.9 |
| | 0.0031 | 0.06 | 0.335 | 107.2 | | 0.0031 | 0.06 | 0.020 | 6.4 |
| | 0.0025 | 0.05 | 0.270 | 108.0 | | 0.0025 | 0.05 | 0.018 | 7.2 |
| | 0.0021 | 0.05 | 0.221 | 106.1 | | 0.0021 | 0.05 | 0.014 | 6.7 |
| | 0.0018 | 0.04 | 0.188 | 105.2 | | 0.0018 | 0.04 | 0.011 | 6.3 |
| | 0.0016 | 0.04 | 0.167 | 106.8 | | 0.0016 | 0.04 | 0.010 | 6.4 |

**Table S3:** Mean inhibition zones [mm] and standard deviation of all 10 API-ILs, 5 ILs and pure ciprofloxacin tested at different concentration per disc.

| | 50 mg/disc | | 25 mg/disc<br>Inhibition zone [mm] | | 10 mg/disc | |
|---|---|---|---|---|---|---|
| | **Mean** | **SD** | **Mean** | **SD** | **Mean** | **SD** |
| Cipro | 29.0 | 0.0 | 25.0 | 0.0 | 18.5 | 0.5 |
| [Chol][CiP] | 26.0 | 3.0 | 21.5 | 2.5 | 14.0 | 3.0 |
| [EMMor][CiP] | 26.0 | 0.0 | 20.5 | 0.5 | 13.5 | 1.5 |
| [TMC$_{10}$A][CiP] | 26.0 | 2.0 | 20.5 | 3.5 | 12.0 | 4.0 |
| [TMC$_{16}$A][CiP] | 26.5 | 2.5 | 21.0 | 4.0 | 15.0 | 4.0 |
| [TC$_8$MA][CiP] | 27.0 | 2.0 | 18.5 | 2.5 | 12.0 | 3.0 |
| [Chol][Cl] | n.d. | n.d. | n.d. | n.d. | n.d. | n.d. |
| [EMMor][Cl] | n.d. | n.d. | n.d. | n.d. | n.d. | n.d. |
| [TMC$_{10}$A][Cl] | 18.0 | 3.0 | 13.0 | 1.0 | 8.5 | 0.5 |
| [TMC$_{16}$A][Cl] | n.d. | n.d. | n.d. | n.d. | n.d. | n.d. |
| [TC$_8$MA][Cl] | 10.0 | 1.0 | 9.0 | 1.0 | 9.0 | 0.0 |

**Table S4:** ½ MIC concentrations used for mutagenesis experiments for all API-ILs, ILs and pure ciprofloxacin investigated in this study.

| API-IL - cation | 1/2 MIC concentration used for Mutagensis experiments [mg/L] | |
|---|---|---|
| | **Cirofloxacin** | **Chloride** |
| Pure antib | 0.01 | |
| [Chol] | 0.02 | 5.0 |
| [EMMor] | 0.03 | 15.0 |
| [TMC$_{10}$A] | 0.03 | 100.0 |
| [TMC$_{16}$A] | 0.03 | 10.0 |
| [TC$_8$MA] | 0.03 | 10.0 |

**Table S5:** Median mutation frequency including 95% confidence intervals and x-fold mutation frequency increase compared to the non-treatment controls for each of the API-ILs, ILs and pure ciprofloxacin.

| | | Median Mutation frequency | Upper Bound 95% CI Range | Lower Bound 95% CI Range | Upper Difference 95% CI Median +/- | Lower Difference 95% CI Median +/- | xfold increase compared to non treatment control |
|---|---|---|---|---|---|---|---|
| **n = 25** 5 technical replicates on 5 different days | Non-treatment | 1.46E-09 | 2.79E-09 | 7.00E-10 | 1.33E-09 | 7.60E-10 | 1.00 |
| | Ciprofloxacin | 2.50E-09 | 6.75E-09 | 1.52E-09 | 4.25E-09 | 9.80E-10 | 1.71 |
| | [Chol][CiP] | 5.97E-09 | 9.09E-09 | 2.19E-09 | 3.12E-09 | 3.78E-09 | 4.09 |
| | [EMMor][CiP] | 3.79E-09 | 5.59E-09 | 2.25E-09 | 1.80E-09 | 1.54E-09 | 2.60 |
| | [$TMC_{10}A$][CiP] | 1.02E-08 | 1.56E-08 | 5.61E-09 | 5.41E-09 | 4.60E-09 | 6.99 |
| | [$TMC_{16}A$][CiP] | 7.33E-09 | 1.32E-08 | 3.14E-09 | 5.84E-09 | 4.19E-09 | 5.02 |
| | [$TC_8MA$][CiP] | 8.62E-09 | 1.72E-08 | 4.76E-09 | 8.54E-09 | 3.86E-09 | 5.90 |
| **n = 25** 5 technical replicates on 5 different days | LB | 2.86E-09 | 3.50E-09 | 1.98E-09 | 6.40E-10 | 8.80E-10 | 1.00 |
| | [Chol][Cl] | 3.45E-09 | 8.30E-09 | 2.20E-09 | 4.85E-09 | 1.25E-09 | 1.21 |
| | [EMMor][Cl] | 2.38E-09 | 4.14E-09 | 1.63E-09 | 1.76E-09 | 7.50E-10 | 0.83 |
| | [TMC10A][Cl] | 4.71E-09 | 6.39E-09 | 2.53E-09 | 1.68E-09 | 2.18E-09 | 1.65 |
| | [TMC16A][Cl] | 1.77E-09 | 2.59E-09 | 1.40E-08 | 8.20E-10 | 3.70E-10 | 0.62 |
| | [TOMA][Cl] | 2.35E-09 | 3.73E-09 | 1.49E-09 | 1.38E-09 | 8.60E-10 | 0.82 |
